# Structural insight into translation initiation of the *λ*cl leaderless mRNA

**DOI:** 10.1101/2023.09.02.556006

**Authors:** Francisco J. Acosta-Reyes, Sayan Bhattacharjee, Max Gottesman, Joachim Frank

## Abstract

In bacteriophage *λ* lysogens, the *λ*cI repressor is encoded by the leaderless transcript (lmRNA) initiated at the *λ*pRM promoter. Translation is enhanced in *rpsB* mutants deficient in ribosomal protein uS2. Although translation initiation of lmRNA is conserved in bacteria, archaea, and eukaryotes, structural insight of a lmRNA translation initiation complex is missing. Here, we use cryo-EM to solve the structures of the uS2-deficient 70S ribosome of host *E. coli* mutant *rpsB11* and the wild-type 70S complex with *λ*cI lmRNA and fmet-tRNA^*fMet*^. Importantly, the uS2-deficient 70S ribosome also lacks protein bS21. The anti-Shine-Dalgarno (aSD) region is structurally supported by bS21, so that the absence of the latter causes the aSD to divert from the normal mRNA exit pathway, easing the exit of lmRNA. A π-stacking interaction between the monitor base A1493 and A(+4) of lmRNA potentially acts as a recognition signal. Coulomb charge flow, along with peristalsis-like dynamics within the mRNA entry channel due to the increased 30S head rotation caused by the absence of uS2, are likely to facilitate the propagation of lmRNA through the ribosome. These findings lay the groundwork for future research on the mechanism of translation and the co-evolution of lmRNA and mRNA that includes the emergence of a defined ribosome-binding site of the transcript.

## Introduction

Translation of genetic information into protein represents one of the most fundamental processes of living organisms. Translation is performed by ribosomes and is divided into four phases: initiation, elongation, termination, and recycling. Despite sharing the same global steps for translation, not all organisms perform translation in the same way, as they have evolved to optimize specific characteristics. Translation initiation is the most divergent and complex process among the different kingdoms of life. It is a highly regulated complex multistep process that requires several initiation factors^1^. It plays a key role in ensuring proper positioning and recognition of the start codon by the initiator tRNA: fmet-tRNA^*fMet*^ and promoting subunit joining to form the 70S elongation-competent complex (70SEC), ready for nascent polypeptide elongation^2^. One of the main differences between translation initiation in eukaryotes and prokaryotes lies in the characteristics of their respective mRNAs. Eukaryotes perform a cap-dependent translation initiation, which involves at least nine initiation factors, fmet-tRNA^*fMet*^ and the messenger-RNA (mRNA), which has specific features such as the 5’-cap, the poly-A tail, the Kozak consensus sequence, and the initiation codon which is recognized during scanning of the mRNA^3,4^. Prokaryotic translation initiation involves three initiation factors and an fmet-tRNA^*fMet*^ as initiator tRNA, but the mRNA is not scanned. Instead, a ribosome binding site (RBS), located in the 5’ UTR region, is utilized for placing the mRNA into the initiating position. The RBS contains the Shine-Dalgarno (SD) sequence, complementary to the 5’-end of the 16S RNA, or anti-Shine-Dalgarno (aSD) sequence^5,6^. This interaction helps position the initiation codon at the P site, allowing proper recognition of the fmet-tRNA^*fMet*^. However, aSD-SD base pairing interaction is not essential for initiation at the correct sites, suggesting that other 5’ mRNA sequences play a role in ribosome alignment as well. In fact, 30 – 50% of *E. coli* mRNAs lack the canonical SD sequence ^7–12^.

An alternative initiation pathway, relevant to this work, has been described for transcripts without a leader, namely leaderless-mRNA (lmRNA), which lacks the upstream 5’ UTR region. These lmRNAs are present in organisms of all kingdoms and are particularly prevalent in *Mycobacterium*, where they represent up to 26% of its genome^13,14^. Translation of lmRNA can follow canonical initiation or the alternative non-canonical initiation^15,16^ in which lmRNA can be directly loaded on the 70S^17^ or 80S^16,18^ ribosome in the presence of the initiator tRNA.

Bacteriophage *λ* has long served as a model organism for studying gene regulatory mechanisms. The *λ*cI repressor is a critical component of the genetic switch that allows the transition of the phage from lytic to lysogenic development^19,20^. The repressor inhibits *λ* transcription and thus permits it to maintain lysogeny. In bacteriophage *λ* lysogens, the *λ*cI repressor is encoded by the leaderless transcript initiated at the *λ* pRM promoter. A down-stream box (DB) sequence (5’-AGCACA-3’) enhances translation of the *λ*cI lmRNA. Host *E. coli* ribosomes lacking ribosomal protein uS2 translate *λ*cI gene more efficiently than wild-type ribosomes (Fig. 1J)^21^. Without the Shine-Dalgarno sequence, translation initiation occurs on the mutant 70S complex rather than the 30S initiation complex. *E. coli* mutants *rpsB11* and *rpsB1* have improved lmRNA translation efficiency. These were characterized and have been the subject of continuous research^15,21–27^. So far however, no structural information is available to explain the improved translation efficiency of the *λ*cI lmRNA by uS2-deficient host ribosome. We selected mutant *rpsB11* for this study. This mutation is an insertion of the transposable element IS1 next to the uS2 stop codon, which reduces cellular levels of uS2. It only limits the availability of uS2 without modifying the protein and results in two types of ribosomes: those with and those without protein uS2 in the cell. We were therefore able to analyze ribosome structures with and without uS2 in the same sample.

**Figure 1.**
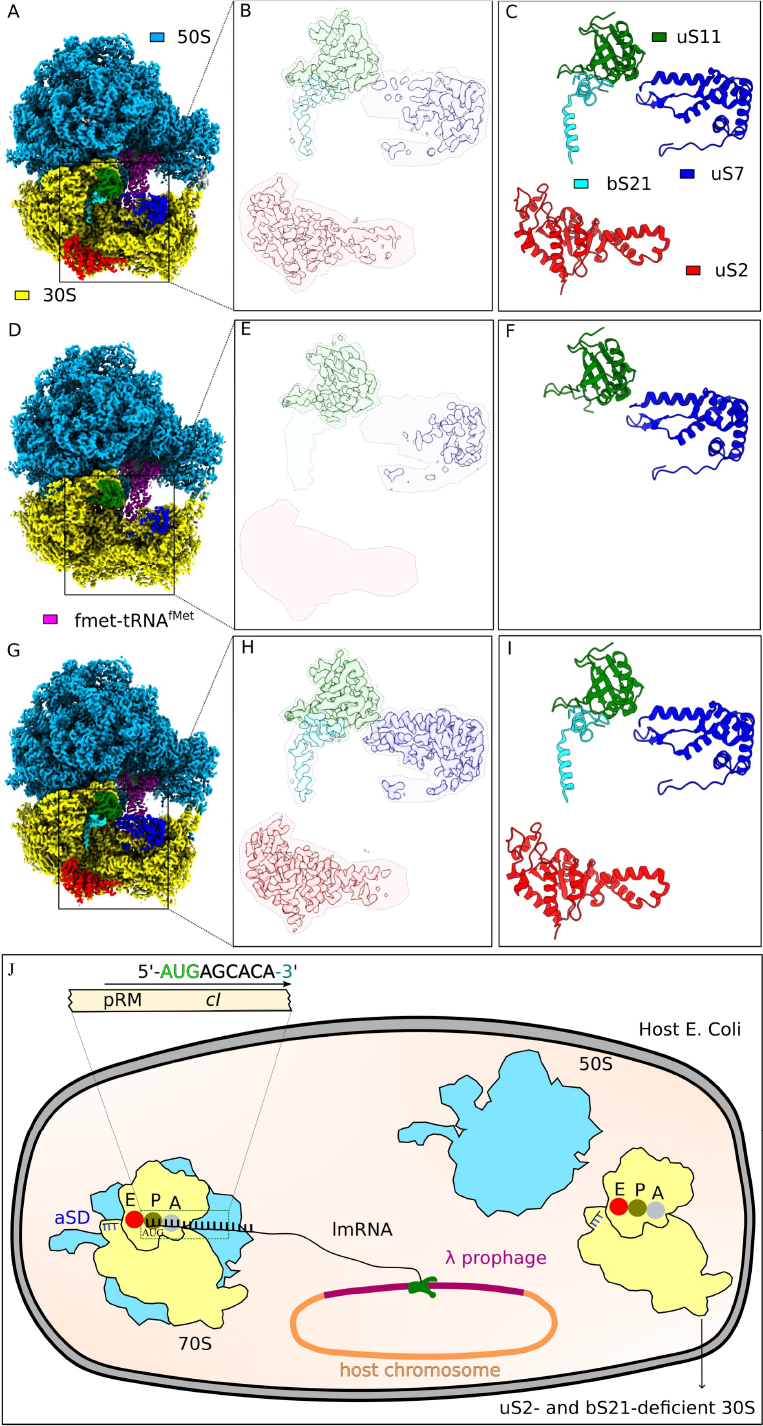
Comparison between ribosomes lacking or containing uS2/bs21 isolated from an *rps*B11 mutant preparation. A) Reconstruction of the wB11. The zoomed view of Coulomb densities and corresponding models of uS2, bS21, uS11, and uS7 proteins are shown in (B) and (C) respectively. (D) is the reconstruction of the nB11. The zoomed view of models of uS2, bS21, uS11, and uS7 proteins are shown in (F), and the Coulomb densities for uS11, and uS7 are shown in (E). The densities of uS2, and bS21 are missing and indicated with their background colors. (G-I) are the same as (A-C), showing MR600. (J) Represents translation initiation of the *λ*cI repressor encoded by lmRNA initiated at the pRM promoter in bacteriophage *λ* lysogens.

In this work, we investigated the role of ribosomal protein uS2 and the structural features that characterize and explain the improved ability of mutant *rpsB11* to translate *λ*cI lmRNA. To this end, we solved the structure of the 70SEC of *E. coli* mutant *rpsB11* loaded with leaderless mRNA and fmet-tRNA^*fMet*^ (B11), using single-particle cryo-EM. The same complex of 70SEC from *E. coli* wild-type strain MRE600 (MR600) was used as control. A minimal *λ*cI lmRNA with 12 bases was used, which comprised the *λ*cI DB sequence (5’-AUGAGCACAAAA-3’) shown to be important in the translation of *λ*cI lmRNA^1^. This *λ*cI lmRNA spans the P site, the A site, and the entry channel without protruding from the ribosome. Our cryo-EM structures reveal that *rpsB11* ribosome contains two ribosome populations: (1) 70SEC without uS2 and uS21 (nB11), and (2) MR600-like 70SEC possessing both uS2 and uS21 (wB11). The uS2-deficient ribosomes not only lack bS1, which binds to the ribosome using uS2 as a scaffold, but are also deficient in bS21. Protein bS21 is normally in contact with the aSD, and its absence repositions the aSD, which would otherwise interfere with the positioning of leaderless transcripts in the mRNA exit channel. We find that in nB11 the absence of bS21 and uS2 increases the 30S head dynamics compared to either MR600 or wB11, which produces charge flow and neutralization through the mRNA entry channel and thereby increases efficiency in translation of *λ*cI lmRNA transcripts.

## Results

### 70SEC structure with a leaderless mRNA

For the cryo-EM study, purified 70S ribosomes from both *rpsB11* and MRE600 strains were mixed *in vitro* with *λ*cI lmRNA and fmet-tRNA^*fMet*^ to produce B11 and MR600 complexes, respectively. Cryo-EM datasets were collected for both complexes (see Methods) and processed following the strategies shown in Fig. S1. After refinement, inspection of the B11 70S consensus map showed poor and broken density in the region of ribosomal protein uS2, indicating a mixed population of both wB11 and nB11, in line with previous result by Shean and Gottesman^21^ that ribosomes from the *rpsB11* mutant are deficient in uS2 protein. Further 3D classification (without re-alignment) resulted in three major classes: two for 70S ribosomes (termed C1m and C2m) and one for the 50S subunit. These classes differ distinctly in the presence (C1m) vs. absence (C2m) of proteins uS2 and bS21 (Fig. 1A-F; Fig. S1). Final refinements on both classes yielded refined structures of nB11 and wB11 with resolutions of 3.3A and 3.2A, respectively (Fig S2). In each complex the fmet-tRNA^*fMet*^ is in the P site, forming a codon-anticodon base pairing with the AUG start codon of *λ*cI lmRNA (Fig. 2A-B).

**Figure 2.**
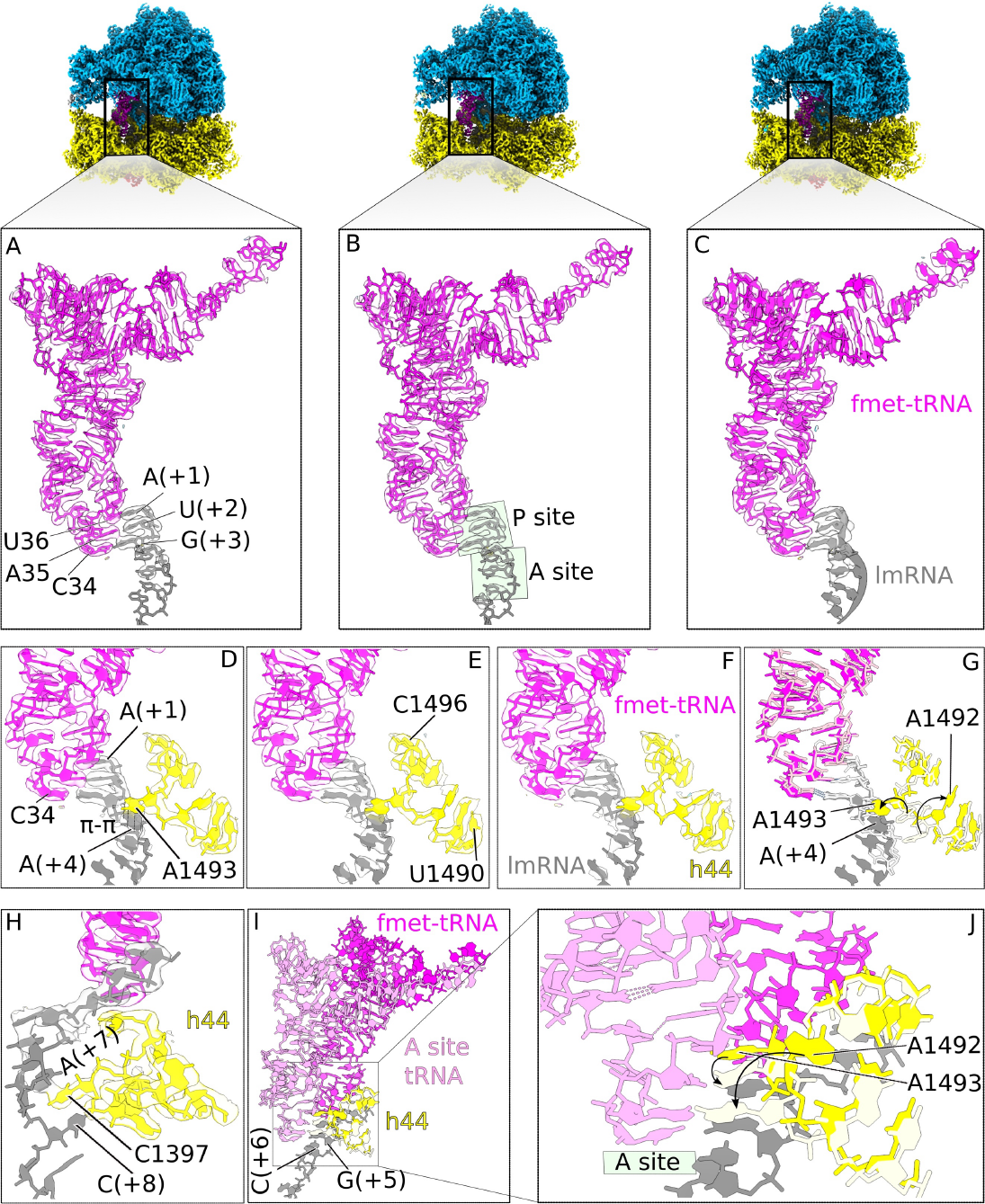
The *λcl* lmRNA interaction with both tRNA and 16S rRNA. (A), (B), and (C) are the *λcl* lmRNA interactions with fmet-tRNA^*fMet*^ for wB11, nB11, and MR600, respectively. The pi-pi stacking between h44 base A1493 with A (+4) of the *λcl* lmRNA for wB11, nB11, and MR600 are shown in (D), (E), and (F), respectively. (G) is the superposition of the leadered mRNA (white) with the *λcl* lmRNA (gray) models. The h44 (yellow) base A1493 flipped to interact with *λcl* lmRNA and consequently, the A1492 base flipped simultaneously from its position (light yellow). (H) is the interaction of (+7) and (+8) bases of*λcl* lmRNA with h44 base C1397 forming a sandwich-like stacking. (I) shows superimposed A-site tRNA (pdb id 4V5D^46^) with P-site initiator tRNA pairing with AUG of *λcl* lmRNA. The sandwich stacking allows *λcl* lmRNA to adopt a conformation suitable for the next upcoming A-site tRNA. (J) is the zoomed view of (I) shows the A-site tRNA needs to replace the A1493 of h44 to form base pairing with the A-site codon and A1492 subsequently flipped back to its original position.

Further focused classification was performed on both wB11 and nB11 using a mask covering the whole region, including bS1, uS2, bS21 and the aSD. After focused classification followed by refinement, some scattered density appears in the region of bS1 in the map of wB11 but not in nB11, evidently since docking of bS1 onto the ribosome requires the presence of uS2. The scattered appearance of bS1 is expected, given the highly flexible nature of this protein. Indeed, there is no single ribosome structure in the pdb with full-length bS1 modeled; those available show only domain 1 (D1) at medium resolution^28,29^. The MR600 complex (Fig. 1G) was processed in the same way as described above for the other complexes, using the same masking region for focused classification. Comparison of reconstructed density maps between MR600 and wB11 shows that the ribosomes are virtually identical (Fig. 1A-C and 1G-I) and in canonical non-rotated configuration, with initiator fmet-tRNA^fMet^ in the P site interacting with the *λ*cI lmRNA start codon AUG (Fig. 2A and C). In both cases the masses of density for protein uS2 as well as bS21 are similar (Fig. 1B-C and 1H-I), and all proteins appear to be in the same configuration when we compute the RMSD among all ribosomal proteins from both MR600 and wB11(Fig. S6). For nB11, although we did not find any densities for proteins uS2 and bS21 (Fig. 1E-F), their absence does not affect the overall ribosomal configuration; nB11 ribosomes remain similar to MR600 (Fig. 1D and G). Both are in the canonical non-rotated configuration, with initiator fmet-tRNA^fMet^ in the P site interacting with the *λ*cI lmRNA (Fig. 2A-B). The shares of populations calculated for nB11 and wB11 -- out of the total number of particles after 3D classification -- are 50% and 10%, respectively.

### Dynamics of the head in the absence of ribosomal protein uS2

Analysis of the refined maps of nB11 and wB11 revealed a noticeable decrease in local resolution in the head of 30S for nB11 compared with wB11 (Fig. 3A-B). Common sources for loss of local resolution can be compositional or conformational; the latter is accentuated in flexible regions. It is known that the motions of the 30S subunit head are essential for canonical translation initiation^30^. In an empty 70S ribosome, the subunit head is stabilized into a single configuration only upon binding of mRNA and P-site tRNA^31^. Although the 70SEC complex is not expected to exhibit substantial movements, the drop in local resolution led us to explore the enhanced dynamics of the 30S head in nB11. For better comparison, Multi-body Refinement^32^ was employed on all three 70SEC complexes: wB11, nB11, and MR600. After analysis of principal components, only the first three components were kept for each complex, as they accounted for almost 50% of the inter-particle variance and stood out in the variance-vs-eigenvalue plot. Most importantly, all three components for nB11 show that the removal of uS2 and bS21 substantially increases the dynamics of the 30S head (Fig. 3C-D). The range in its angle of rotation almost doubles as compared to the other two complexes (5.6^0^ for wB11 and 10.6^0^ for nB11). We also observed the “nodding” latch-opening/closing head movement seen for small subunits at the mRNA entry channel^33^ (Fig. S7). For wB11 and MR600, we observe normal ratchet-like motions (Fig. 3D) along with the latch opening.

**Figure 3.**
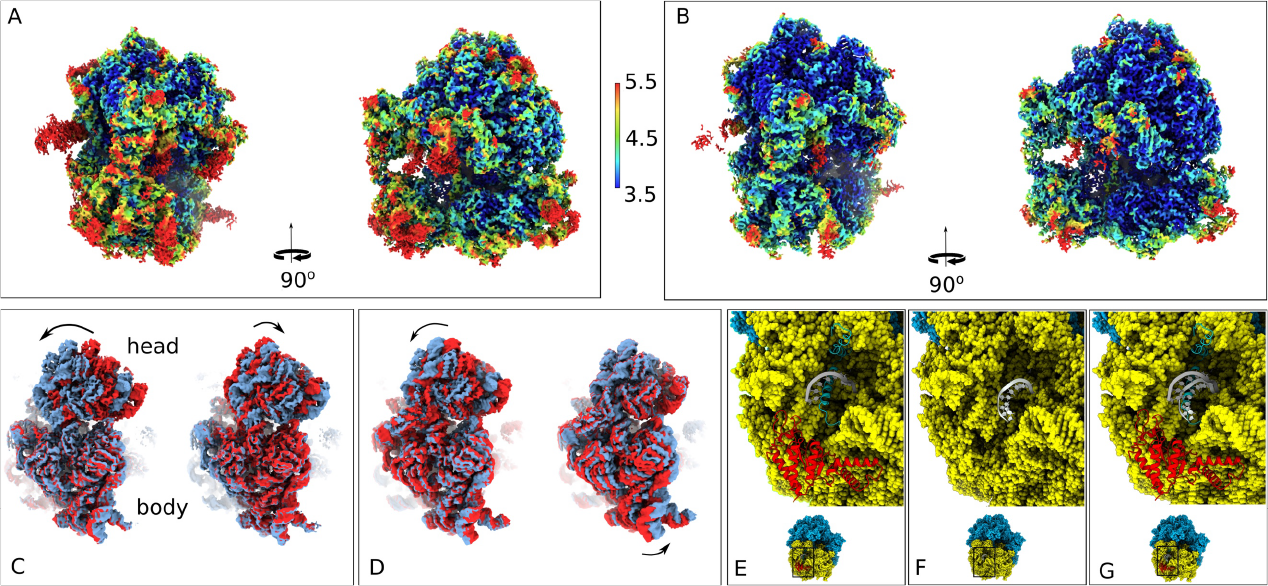
Dynamics of nB11 30S head and aSD shift. Local resolution estimation of (A) nB11 and (B) wB11. The first and end states of the first two principal components are compared to show the increased 30S head dynamics of (C) nB11 compared to the same for (D) wB11. The support to aSD (gray) provided by bS21 (sky blue) is shown in (E) in zoomed view of the wB11 corresponding region. The lack of bS21 (cyan) leads to the transition of aSD (light gray) is shown in (F) as a zoomed view of nB11 corresponding region and its superimposition with wB11 to show the noticeable displacement of aSD clearly and is shown (G).

The unusual motions detected by Multi-body Refinement as well as the drop in local resolution for the 30S subunit in nB11 prompted us to do focused classification using a mask including the 30S subunit, which resulted in three distinct classes of the 30S subunit, which we termed C3m (42%, 44,212 particles), C4m (8%, 8,422 particles), and C5m (33%, 34,737 particles) (Fig S1). Subsequent refinements followed by atomic model building were performed on all three classes (Fig 4A-C, S1, and S5). In class C3m, fmet-tRNA^*fMet*^ is in the classical P site, with the 30S head un-swiveled and the body in the non-rotated state (Fig. 4A). In class C5m, fmet-tRNA^*fMet*^ has moved from the P site to the intermediate hybrid pe/E state, between P and E site, as a consequence of 30S head swiveling (by 12.1^0^) and 30S body rotation (by 9.7^0^) (Fig. 4B-C, and 4F-H, S4). In class C4m we observe the 30S head slightly swiveled (1.5^0^) and 30S body slightly rotated (2.4^0^) to an intermediate state between C3m and C5m, and the fmet-tRNA^*fMet*^ situated at the P site as in class C3m (Fig. S5A-F). We repeated the same focused classification step for wB11 as well as MR600 but failed to find any subpopulations in this case.

**Figure 4.**
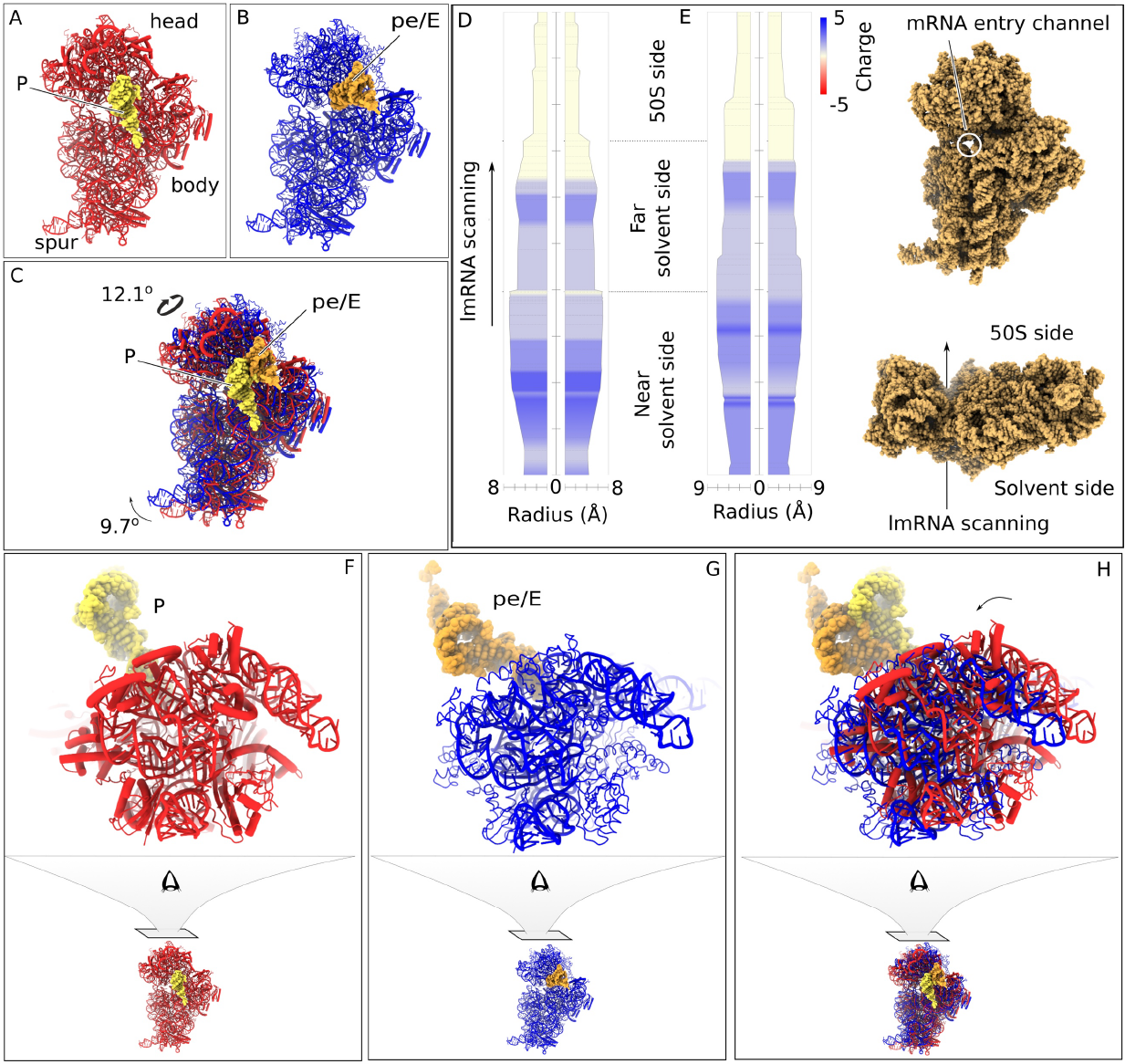
Dynamics and charge flow in the mRNA entrance channel due to 30S head and body movement in nB11. (A), atomic model of C3m (red) with the P site fmet-tRNA^*fMet*^ (yellow). (B), atomic model of C5m (blue) with pe/E site fmet-tRNA^*fMet*^ (light-brown). (C), superimpositions of C3m and C5m showing the movement of fmet-tRNA^*fMet*^ from the P site to the hybrid pe/E state. (D) and (E), variation of radius and Coulomb charge along the *λ* cl lmRNA channels of classes C3m and C5m, respectively. The averaged charge distribution in the channel is shown in red (-charge), blue (+ charge), and yellow (neutral) color. The position of the *λ* cl lmRNA channel and the direction of *λ* cl lmRNA propagation are shown in the side panel. (F-H), zoomed view of the head swiveling of the 30S subunit between C3m and C5m.

### The enhanced head dynamics causes Coulomb charge flow through the mRNA entry channel

The different 30S subunit states and head rotations of nB11 go hand in hand with conformational differences in the mRNA entry channel, which are expected to affect the movement of *λ*cI lmRNA through the channel. Following this line of reasoning, we compared the dynamics and charge distributions of the mRNA entry channel for the three atomic models of the C3m, C4m, and C5m, by measuring the channel diameter and Coulomb charge distributions of the constituting residues in a similar way as done for membrane proteins (see Methods). Interestingly, we found notable differences in both radius and charge distribution (Fig. 4D-E). The residues responsible for these differences belong to 16S rRNA and to the near-solvent site 30S subunit proteins uS3, uS4, and uS5. Comparison of channel radius between the two 30S subunit models of predominant classes C3m and C5m reveals a peristalsis-like movement from the near-solvent side to the 50S subunit side, or in the direction of the required *λ*cI lmRNA scanning (Fig. 4D-E, S10, movie 1). The mRNA entry channel diameter increases on the 50S subunit side, while it decreases on the near-solvent side when the 30S head swivels from class C3m to C5m. In summary, the behavior can be described as a flow of positive Coulomb charges from the near-solvent side toward the 50S subunit side, and a charge neutralization at the far solvent side of the mRNA entry channel when going from the C3m to the C5m state (Fig. 4D-E, S10). The overall charges are positive at the near-solvent side, as they are formed by positively charged amino acids of the proteins uS3, uS4, and uS5 (Fig. 4D-E) while they remain neutral at the 50S subunit side, where codon anti-codon base pairing between the *λ*cI lmRNA and tRNA occurs. Class C4m, in an intermediate state of head swiveling and body rotation between C3m and C5m, does not present an intermediate in charge distribution but only represents a minor population (Fig. S5G).

### The configuration of *λ*cI lmRNA and its interaction with the initiator tRNA

The leaderless transcript is loaded in a configuration identical in both 70SEC classes C1m and C2m, i.e., the presence or absence of uS2 and bS21 has apparently no impact on the conformation of *λ*cI lmRNA (Fig 2A-F). Nevertheless, mRNA bases downstream toward the vacant A site (+4 to +8), following the start codon, are stabilized to a conformation suitable to form codon-anti-codon base pairing with upcoming A-site tRNA, through stacking interaction with h44 of 16S rRNA (2H-I). The downstream *λ*cI lmRNA base A_+4_ is found to form a π-stacking interaction with the monitor base A1493 of h44 (Fig. 2D), while C1397 of h44 is intercalated between A_+7_ and C_+8_ (Fig. 2H). However, the monitor base A1943 of h44 needs to be flipped to accommodate A-site tRNA (Fig. 2G).

### The lack of uS2/bS21 affects the structure and properties of the mRNA exit channel

The increased head dynamics of nB11 in the absence of proteins uS2/bS21 affect not only the mRNA entry channel but also those of the mRNA exit channel, by destabilizing the interaction between proteins uS7 and uS11. Density in the region of the *β*-hairpin of uS7 appears fragmented (Fig. 1D-E). This behavior agrees with the spatial distribution of the B-factor: a higher B-factor is evident in the region of the interaction between uS7 and uS11, most notably in the *β*-hairpin of uS7 (Fig. S8A-B). The RMSD of the superimposed atomic models of uS7 and uS11 extracted from MR600 and nB11 shows a noticeable deviation at the *β*-hairpin of uS7, in agreement with the largest (11Å) distance between the *β*-hairpin of uS7 and uS11 (Fig. S8C). Notably, for both complexes, upon focused classification on the whole region around bS1, uS2, bS21, and aSD, we observe differences in the position of the aSD (Fig. 3E-G). Although the densities corresponding to the aSDs were not resolved with high resolution, the differences in the aSD position are clearly evident (Fig. S9).

## Discussion

### Impact of the missing 30S proteins uS2 and bS21 in the *rpsB11* mutant

The classification revealed that *rpsB11* possesses a major population of nB11 ribosomes in which both uS2 and bS21 proteins are missing. On the other hand, the map of wB11, comparatively less populated, does contain both proteins, in agreement with the fact that in *rpsB11* the mutation reduces the production of protein uS2. The wB11 complex proves to be identical to the MR600, and thus represents the minimal fraction in the number of ribosomes that are preserved by the cell for efficient leadered mRNA translation following the canonical SD-aSD recognition mechanism (Fig. 5A-B). The 30S subunit assembly map reveals that both uS2 and bS21 are tertiary binding proteins, as they are incorporated into the ribosome in the later stages of assembly and require at least one protein from each of the primary (such as uS4, uS7) and secondary (such as uS5, uS11) bonding proteins for their stable assembly^35^. We noticed that in *rpsB11* protein bS21 is missing whenever uS2 is missing. This can be explained by the enhanced dynamic nature of both uS11 and uS7 due to the increased head dynamics of uS2-depleted ribosomes. This tends to disrupt the stable architecture formed by the 30S head, platform, uS11, and uS7, which is necessary for the stable binding of bS21 and its downstream functions (Fig 1G-I).

**Figure 5.**
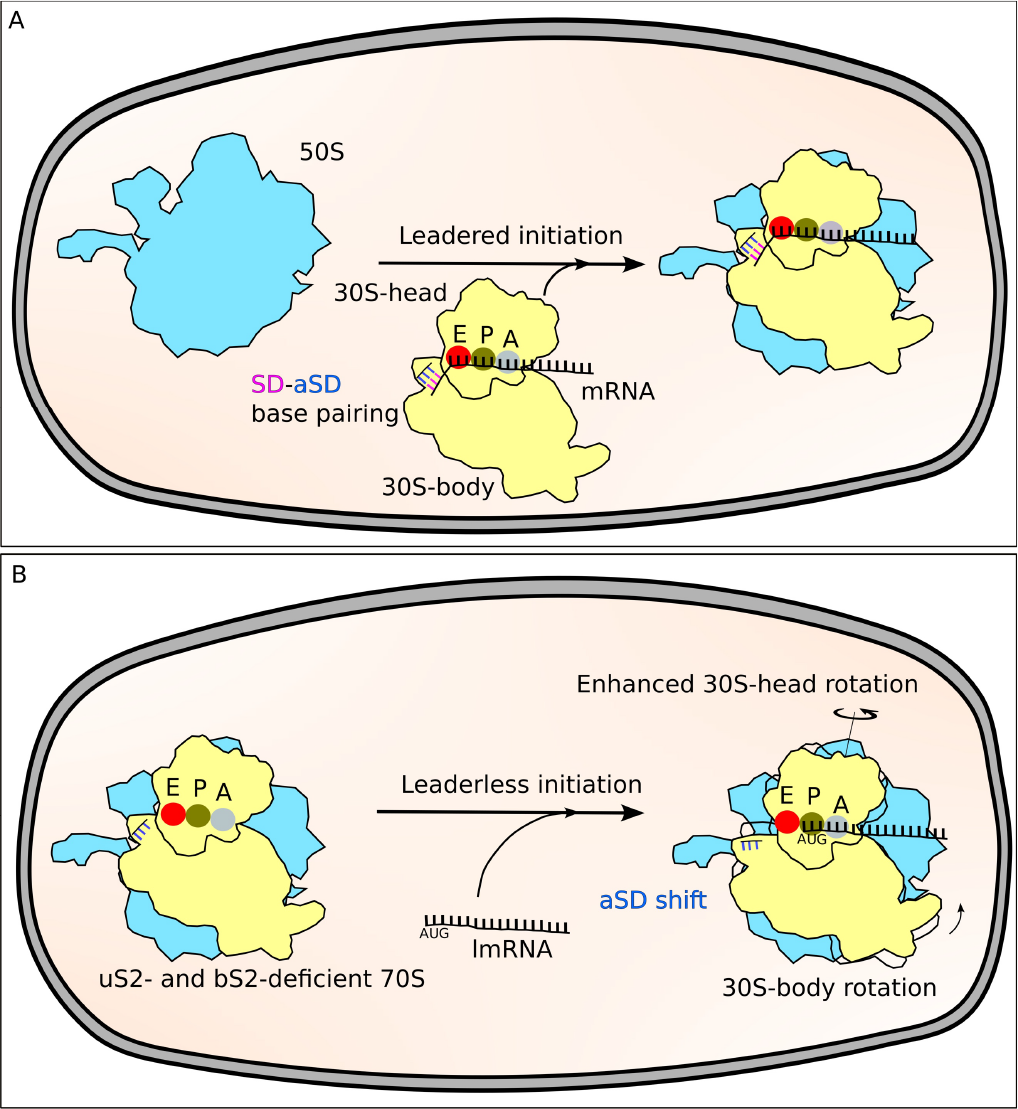
A comparison of SD-led and leaderless mRNA translation initiation. The formation of the 70S initiation complex (A) is represented by the association of the 50S with the 30S connected with leadered mRNA via base pairing between SD and aSD. (B) shows the direct interaction of leaderless mRNA with the uS2- and bS21-deficient 70S to begin translation. Additionally, the improved head dynamics of mutant 70S aids in the threading of lmRNA and the elongation of the nascent polypeptide chain. In this representation, the positions of tRNAs and other initiation factors are concealed.

As reported, ribosomes that lack bS21 or have been treated with antibodies against bS21 can perform the majority of reactions of unmodified ribosomes^36^. The enhanced head dynamics of nB11, which lacks bS21, can explain this fact, as it affects the architecture of both mRNA entrance and exit channels. Both changes are described below.

In the first case, a peristalsis-like motion is observed in the mRNA entrance channel of nB11, facilitated by the increased head swiveling of the 30S subunit. When the 30S head swivels from the non-rotated to a rotated state, the mRNA entry channel diameter increases on the 50S subunit side, whereas it decreases on the near-solvent side. This combination of changes is likely to promote the threading of *λ*cI lmRNA through the channel (Fig. 4D-E, movie-1). Along with the change in channel diameter, we not only notice the flow of positive Coulomb charges through the channel from the near-solvent toward the 50S subunit side, but also a charge neutralization. The 30S head rotation exposes side-chain atoms with different charges and polarities, resulting in charge flow and neutralization inside the mRNA entry channel. The flow of positive Coulomb charge may be instrumental in the translocation of the negatively charged *λ*cI lmRNA (Fig 4D-E, movie-1). The additional neutralization during swiveling of the head to the rotated state may accelerate the movement of negatively charged *λ*cI lmRNA through the mRNA entry channel. Interestingly, in both classes, the mRNA entry channel at the near-solvent side is positively charged because of the presence of positively charged constituent amino acids of the proteins uS3, uS4, and uS5. Despite the presence of negatively charged 16S rRNA, the appearance of positive charge is expected to help negatively charged *λ*cI lmRNA in enabling entry of their 5’ end into the channel. Additionally, it is likely that the neutral charge distribution at the 50S subunit side will not compensate for the *λ*cI lmRNA charge needed for the codon-anticodon base pairing between *λ*cI lmRNA and tRNA. It is important to note that the same mechanism will hold for leadered mRNA, but the effect would be smaller, as pointed out above, because of the smaller range of the head swiveling.

In the second case, the aSD is shifted away from the path of the mRNA emerging from the mRNA exit channel. It is well known that SD-aSD interaction promotes the recognition of leadered mRNA^37^. Although the aSD adopts a shifted position in the absence of bS21, it is not constrained in this position, and SD recognition can still occur with the remaining leadered mRNAs available within the cell. Comparison of atomic models at aSD regions of nB11 and MR600 revealed that bS21 acts as a support to hold the flexible aSD. In the case of nB11, no bS21 is present and, due to the lack of support, and the aSD has shifted from its original position (Fig 3E-G, S9). It has been proposed that bS21 interacts with nearby mRNA sequences 5’ of the Shine-Dalgarno helix, thereby affecting translation initiation^37,38^

The RNA exit channel is constituted of the 30S head and platform and is stabilized by the interaction between uS7 and uS11. Mutations of residues involved in this interaction are known to impact translation initiation, affecting the interaction between the 30S subunit and mRNA, as well as translational fidelity^39^. Disruption of this interaction, as suggested, will result in an increase in head dynamics. For ribosomes in nB11, however, the converse appears to be true. Here, the increased dynamics of the head caused by the absence of uS2/bS21 (Fig. 1D and 3C) has apparently resulted in disruption of the interaction between uS7 and uS11 (Fig. 1E). The *β*-hairpin loop of uS7 acts as a sort of gate behind the E site, where a substantial kink of the mRNA occurs, in part guided by the *β*-hairpin loop. Initiation factor IF3 binds near this region, and, together with the proper configuration of the head, imposes an open-latch conformation on the 30S subunit^33^. The motion of uS7 along with the 30S subunit head facilitates mRNA sliding. This explains why disruption of the contact between uS7 and uS11 increases frameshifting, stop codon readthrough, as well as misreading^39^. In the case of leaderless transcripts, this disruption results in widening the exit channel, potentially facilitating the threading of *λ*cI lmRNA through the channel.

### Stabilization of the *λ*cI lmRNA by 16S rRNA facilitates stable fmet-tRNA^*fMet*^ interaction

The *λ*cI lmRNA was found to adopt a stable conformation when binding with fmet-tRNA^*fMet*^. This stable conformation is facilitated by the π-stacking interactions between h44 of 16S rRNA and *λ*cI lmRNA bases A_+4_ and A_+8_. Although this conformation makes the downstream bases A_+4_ to A_+6_ easily available for the next upcoming A-site tRNA (Fig. 2I), the monitor base A1493 of helix h44 needs to be flipped for compatibility with the anticodon of A-site tRNA (Fig. 2I-J). A similar configuration has been identified recently in a eukaryotic ribosome with a poly-A mRNA, where the poly-A tract interacts with A1825 and C1698^40^. We suggest that the sequence of bases A_+4_ to A_+7_ is relevant for the stability of this downstream motif. Importantly, position +4 is substantially enriched in adenines for annotated *E. coli* start sites. The positioning of the *λ*cI lmRNA through the mRNA entry channel agrees with previous structures observed^41^, implying that non-specific interactions of arginines (ARG-19 from uS5, ARG-130&163 from uS3, and ARG-46 from uS4) with the *λ*cI lmRNA can also contribute to its stabilization. The fact that these interactions are not specific to the absence of proteins uS2/bS21 suggests that their contribution becomes relevant for translation initiation and correct mRNA stabilization only in the absence of SD interaction.

### The conformational mobility of 70S ribosome lacking uS2 and bS21 can facilitate translocation of *λ*cI lmRNA

During elongation, Elongation Factor G (EF-G) helps to translocate the classical A and P-site tRNAs to the hybrid ap/P and pe/E states, respectively. Simultaneously, the 30S head swivels to a rotated state and the 30S body undergoes ratchet-like rotation. This rotation is crucial for the mRNA frame reading and downstream production of the nascent polypeptide chain^42^. In the absence of EF-G, this hybrid state can also exist, but it requires a lower Mg^2+^ concentration (∼5 mM) than is normally used (10-15 mM) for in vitro translational assays. In the nB11 complex, the P-site fmet-tRNA^*fMet*^ is in the position of the hybrid pe/E state which, along with the 30S head and body rotation, is similar to that is seen for the EF-G-mediated translocation. We observed this state of the nB11 complex at 10 mM salt concentration, so it is clear that this state is not a consequence of a lowered Mg^2+^ concentration. This implies that a 70S ribosome lacking uS2 and bS21 has a larger propensity for factor-independent translocation than wt ribosomes, a possibility that will require experimental support. Although we did not use any tRNA at the A site in our study, it is obvious that if it were present it would shift to the ap/P hybrid state concurrently with the translocation of *λ*cI lmRNA.

### Antibiotic and inhibitor resistance

As we have seen, the increased 30S subunit head dynamics disrupts the architecture of the mRNA exit channel and thereby destabilizes the uS7-uS11 interaction. As the mRNA exit channel is the binding site of antibiotics like Pactamycin and Kasugamycin, the increased head dynamics may inhibit binding of these antibiotics, consistent with the resistance to these drugs by the *rpsB* mutants. Pactamycin interacts with residues G693 and C795 near the mRNA exit channel and inhibits translocation by blocking the E site^43^. Kasugamycin interacts at the E site and at the mRNA exit channel. By mimicking mRNA nucleotides, Kasugamycin thereby destabilizes tRNA binding and disrupts canonical translation initiation^44^. Both Kasugamycin and Pactamycin may enhance the translation of *λ*cI by inhibiting either uS2 itself or ribosomes that contain uS2. In this fashion, these antibiotics would reproduce the phenotype of *rpsB* mutants that overexpress *λ* cI.

Not every bacterium makes use of SD motifs to initiate translational start, and a genome-wide analysis demonstrates that whereas SD motifs promote initiation efficiency, they are not required for ribosomes to select the mRNA location of initiation^8^. So far, uncertainty surrounds the mechanism by which A-rich sequences improve initiation. It was proposed that the A-bases that are bound inside the ribosome and are near the start codon may selectively interact with ribosomal components^8^.

In conclusion, we have revealed the mechanism of leaderless mRNA translation initiation, in which the 30S body stabilizes the *λ*cI lmRNA for base-pairing with fmet-tRNAfMet while the enhanced 30S head dynamics enable *λ*cI lmRNA entry, propagation, and exit through the mRNA entry and exit channels. Moreover, the increased head dynamics can mimic translocation steps to make the process more effective.

Our study thus bears on the underlying regulation of lysogeny but also provides an insight into leaderless mRNA translation, a process that is conserved for bacteria, archaea, and eukaryotes

## Acknowledgements

This work was supported by a grant from the National Institutes of Health P01CA174653 (to M.E.G), and R01GM29169, R35GM139453 (to J.F.). All data were collected at the Columbia University Cryo-Electron Microscopy Center (CEC). We thank Robert A. Grassucci, Zhening Zhang, and Yen-Hong Kao for their help with the cryo-EM data collection.

## Author contributions

M.E.G., and J.F. conceived the research; F.J.A.R., S.B., M.E.G., and J.F. designed the project; F.J.A.R. prepared the biological samples; F.J.A.R., and S.B. processed the cryo-EM data; F.J.A.R., and S.B. did the model building and validation. All authors contributed to the article writing.

## Data and code availability

The refined maps are deposited on EMDB and corresponding atomic models on PDB and will be publicly available as of the date of publication.

**EMD**-41049 (wB11), 41050 (nB11),

**PDB**-8T5D (wB11), and 8T5H (nB11).

## Notes

### Competing Interest Statement

The authors have declared no competing interest.

### Summary of Updates

In bacteriophage λ lysogens, the λcI repressor is encoded by the leaderless transcript (lmRNA) initiated at the λpRM promoter. Translation is enhanced in rpsB mutants deficient in ribosomal protein uS2. Although translation initiation of lmRNA is conserved in bacteria, archaea, and eukaryotes, structural insight of a lmRNA translation initiation complex is missing. Here, we use cryo-EM to solve the structures of the uS2-deficient 70S ribosome of host E. coli mutant rpsB11 and the wild-type 70S complex with λcI lmRNA and fmet-tRNAfMet. Importantly, the uS2-deficient 70S ribosome also lacks protein bS21. The anti-Shine-Dalgarno (aSD) region is structurally supported by bS21, so that the absence of the latter causes the aSD to divert from the normal mRNA exit pathway, easing the exit of lmRNA. A π-stacking interaction between the monitor base A1493 and A(+4) of lmRNA potentially acts as a recognition signal. Coulomb charge flow, along with peristalsis-like dynamics within the mRNA entry channel due to the increased 30S head rotation caused by the absence of uS2, are likely to facilitate the propagation of lmRNA through the ribosome. These findings lay the groundwork for future research on the mechanism of translation and the co-evolution of lmRNA and mRNA that includes the emergence of a defined ribosome-binding site of the transcript.

