## Supplementary material for "Structural insight into translation initiation of the *λ*cl leaderless mRNA": SI

### Methods

#### Purification of ribosomes and transcript description

Ribosomes were purified from the *E. coli* strain *rspB11* and from wild-type MRE600 (as a control). Cells were harvested in the mid-log phase (around OD600 of 0.6) and resuspended in 40 mM Tris-HCl at pH 7.5, 500mM $NH_4Cl$, 10 mM $MgCl_2$, 2 mM DTT, protease inhibitors, and RNase inhibitors. Cell pellets were disrupted by sonication. The lysate was clarified by centrifugation for 20 min at 18,600 g. Ribosomes in the supernatant were pelleted through a 1 M sucrose cushion in the same buffer for 6 hrs. at 44,000 rpm in a Ti45 rotor (Beckman Coulter, Brea, CA). The pellets were resuspended in 20 mM Tris-HCl at pH 7.5, 50 mM KCl, 10 mM $MgCl_2$, 2 mM DTT, and incubated for 5 min with 1 mM puromycin on ice. The sample was loaded on a 10%–40% sucrose gradient in resuspension buffer (20 mM Tris-HCl at pH 7.5, 50 mM KCl, 10 mM $MgCl_2$, 2 mM DTT) and centrifuged for 15 hr at 22,000 rpm in an SW 28 Ti Swinging (Beckman Coulter). For subunit purification, 70S ribosomes were exchanged into dissociation buffer (20 mM Tris-HCl at pH 7.5, 300 mM KCl, 1 mM $MgCl_2$, 2 mM DTT) before loading onto a 10%-35% sucrose gradient in the same buffer and centrifuged for 17 hrs. at 22,000 rpm in the SW 28 Ti rotor. The 50S and 30S subunits were exchanged separately into reassociation buffer (20 mM Tris-HCl at pH 7.5, 50 mM KCl, 10 mM $MgCl_2$, 2 mM DTT), concentrated to 11 μM, aliquoted and stored at -80°C after being flash frozen in liquid nitrogen. Charged fmet-tRNAfMet was kindly donated by Dr. Jingji Zhang. The mRNA oligonucleotide of 12 bases length, which includes the initiation codon AUG and the DB sequence reported by Shean and Gottesman[1], 5'-AUGAGCACAAAA-3', was purchased from Sigma-Aldrich (HPLC purified for crystallographic purposes).

#### Assembly of 70S complex and data collection

Components of 70SEC complexes were mixed and incubated for 20 mins at 37°C for a final concentration of 500 nM 70S, 1μM fmet-tRNAfMet, 2 μM mRNA in Polymix-Buffer. Aliquots of 3μl of assembled ribosomal complexes were incubated for 3 seconds on plasma-treated holey gold grids[2]. Grids were blotted for 5-7s and flash-cooled in liquid ethane using a Vitrobot (Thermo Fisher Scientific, Oregon, USA). Screening of the sample to optimize concentration and grid preparation conditions were carried out in an F20 Tecnai microscope (Thermo Fisher Scientific) operated at 200kV and equipped with Gatan K2 Summit direct detector (Gatan, CA, USA). Datasets were collected in a Polara-G2 microscope (Thermo Fisher Scientific), operated at 300 kV, and equipped with a Gatan K2 direct detector. Data for rspB11 were recorded in super-resolution counting mode at a magnification of 23,000, corresponding to a calibrated pixel size of 0.83 Å (1.66 Å per physical pixel). The WT control was collected with counting mode at a magnification of 39,000, corresponding to a calibrated pixel size of 0.98 Å. Defocus values ranged from 0.8 to 3 μm. Images were recorded in automatic mode using the Leginon[3] software and frames were aligned with Motioncor2[4] and checked on the fly using APPION[5].

**Image processing and structure determination.**

A flowchart of the data processing steps is shown in Fig. S1. 3452 good micrographs were selected for the B11 strain for further data processing. The beam-induced motion of the sample was corrected using the MotionCor2 program. The contrast transfer function (CTF) of each micrograph was estimated using CTFFIND4[6]. Particle picking was performed using Topaz[7]. Good particles were selected by 2D classification and trained using 10,000 particles. Autopicking using the trained Topaz model yielded 609,632 particles. Particles picked by Topaz were subjected to 2D classification for further selection of good particles, which yielded 572,744 particles. All particles were pooled together and used for 3D initial model generation followed by 3D auto-refinement, applying C1 symmetry in RELION4[8]. CTF refinements were done to correct for magnification anisotropy, fourth-order aberrations, per-particle defocus, and per-particle astigmatism, followed by another 3D auto-refinement. Then 3D classification was performed without alignment, using the angular information from the previous refinement step. The 3D classification produced two distinct classes of the 70S ribosome, which differ distinctly in the presence (C1m, 17,544 particles) vs. absence (C2m, 105,266 particles) of proteins uS2 and bS21, and one class of 50S. The 50S class was not included for further structural studies. Particles from each class were further subjected to 3D auto-refinement, and the post-processing was done with RELION4 and DeepEMhancer[9]. Further focused refinements without alignment were done on the 30S subunit part of the map for both C1m and C2m, which resulted in three distinct subclasses for C2m, namely C3m (non-rotated, 44,212), C4m (intermediate, 8,422), and C5m (rotated, 34,737). We did not find any subclasses for C1m, which showed the 30S subunit in a non-rotated position. Local resolutions were estimated using ResMap[10]. A similar approach was applied to MR600. To build models for the 70S ribosomes, pdb id:6XE0[11] for 30S, pdb id:6XZ7[12] for 50S, and pdb id:6O7K[13] for tRNA were selected as starting models. Models were first fitted as rigid bodies using Chimerax1.5[14] and then refined using Phenix real-space refinement[15]. The lmRNA model was built manually in COOT[16]. Further model validations were done using Phenix Comprehensive Validation (cryo-EM) and tabulated in Table S1. Charge and radius distribution of mRNA entrance channel insides were calculated using MOLEonline[17].

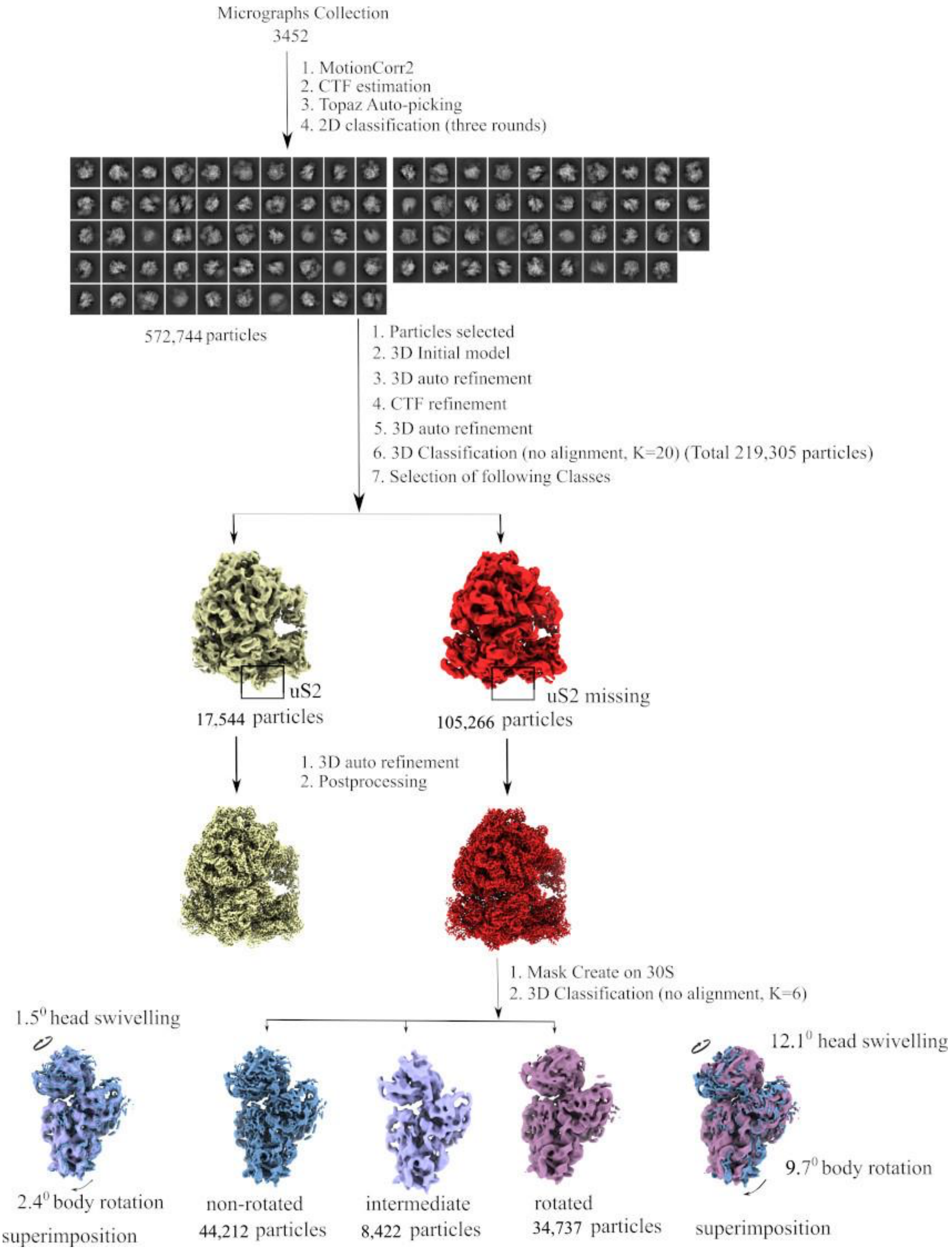


**Figure S1. Workflow of cryo-EM data processing.**

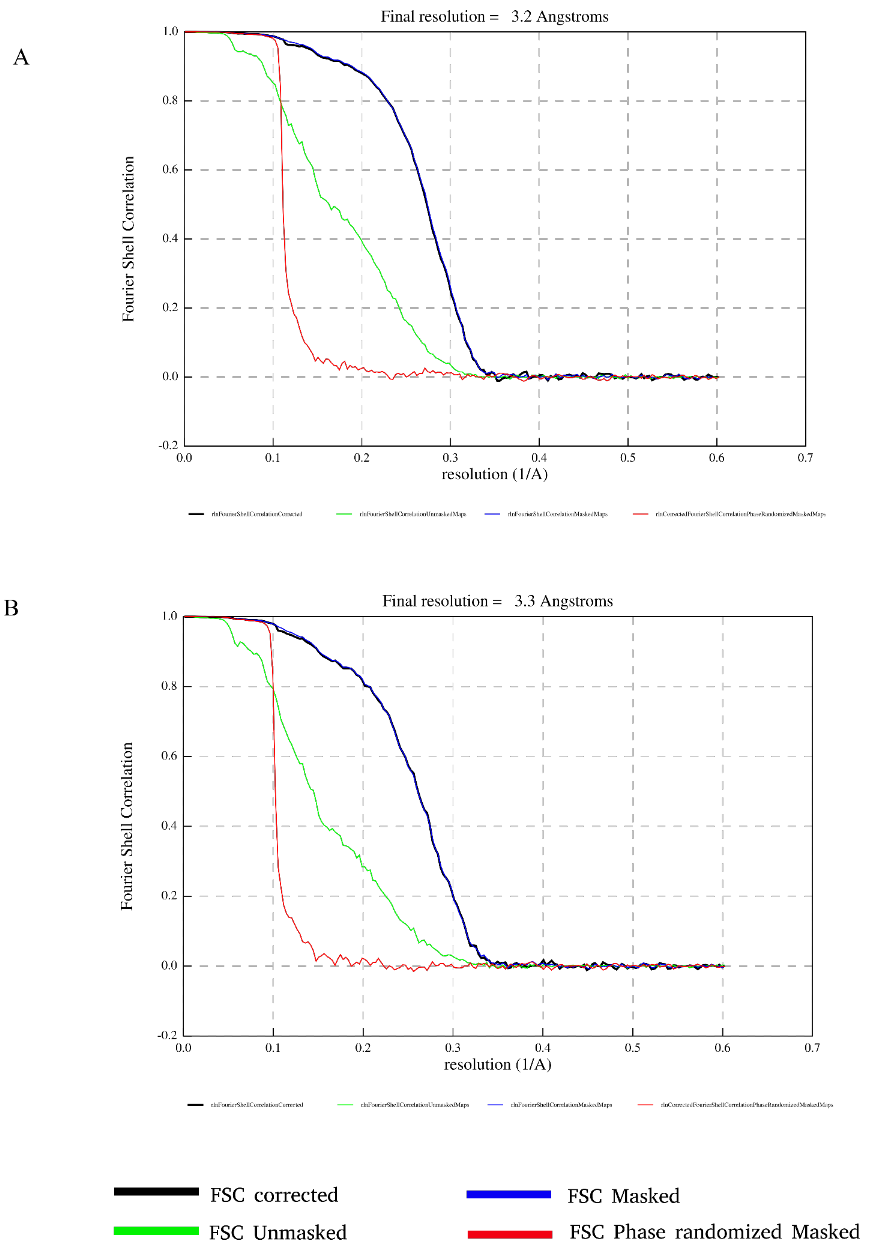


**Figure S2. Resolution estimation.** FSC plots and estimated resolutions of (A) wB11, and (B) nB11ontained from Relion4.

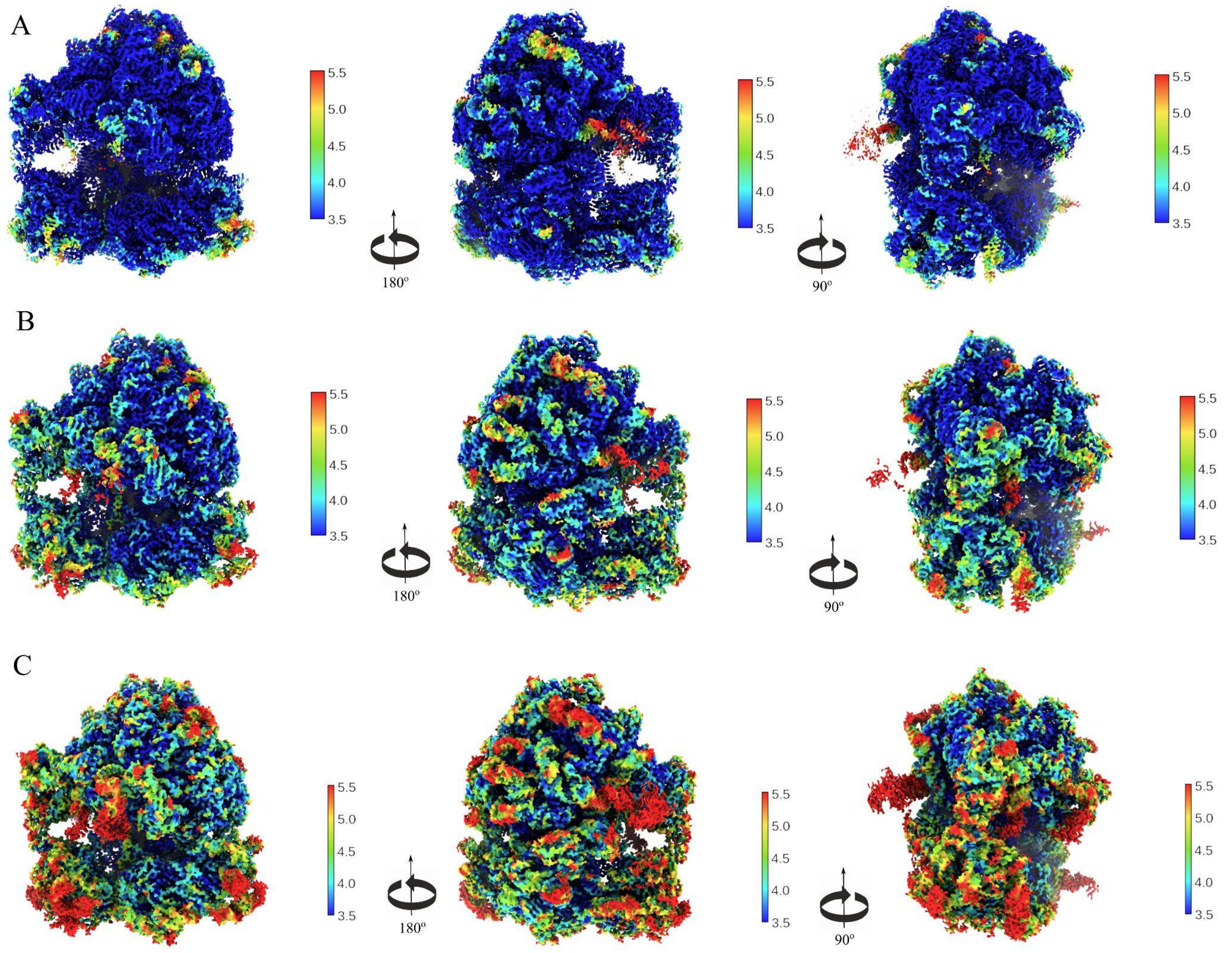


**Figure S3. Changes in local resolution.** Local resolution estimates for (A) MR600, (B) wB11, and (C) nB11.

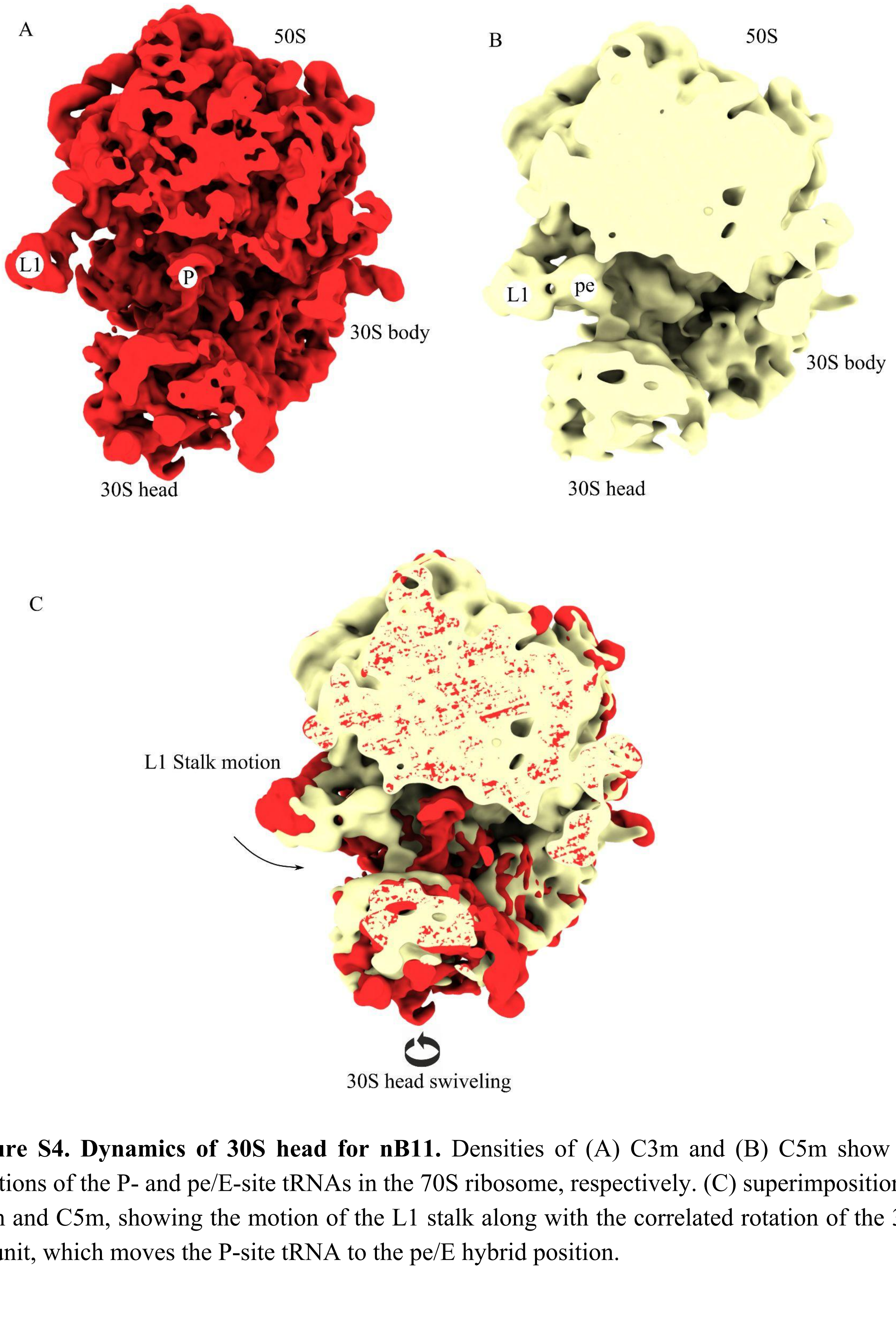


**Figure S4. Dynamics of 30S head for nB11.** Densities of (A) C3m and (B) C5m show the positions of the P- and pe/E-site tRNAs in the 70S ribosome, respectively. (C) superimposition of C3m and C5m, showing the motion of the L1 stalk along with the correlated rotation of the 30S subunit, which moves the P-site tRNA to the pe/E hybrid position.

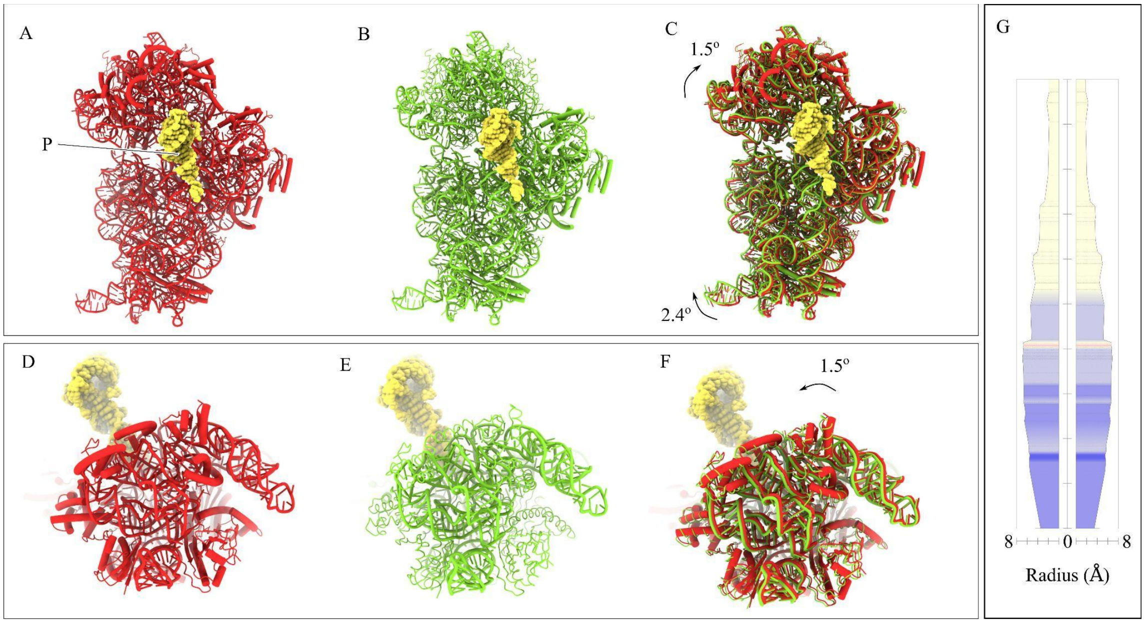


**Figure S5. The lmRNA channel charge distribution due to 30S head and body movement in an intermediate position (supported by small C4m subpopulation only).** (A,B) Atomic models of 30S subunit in C3m and C4m, respectively. (C), superimposition of atomic models in (A) and (B). (D-F) 30S head swiveling from C3m (red) to C4m (green). (G), charge distribution in the lmRNA channel for C4m.

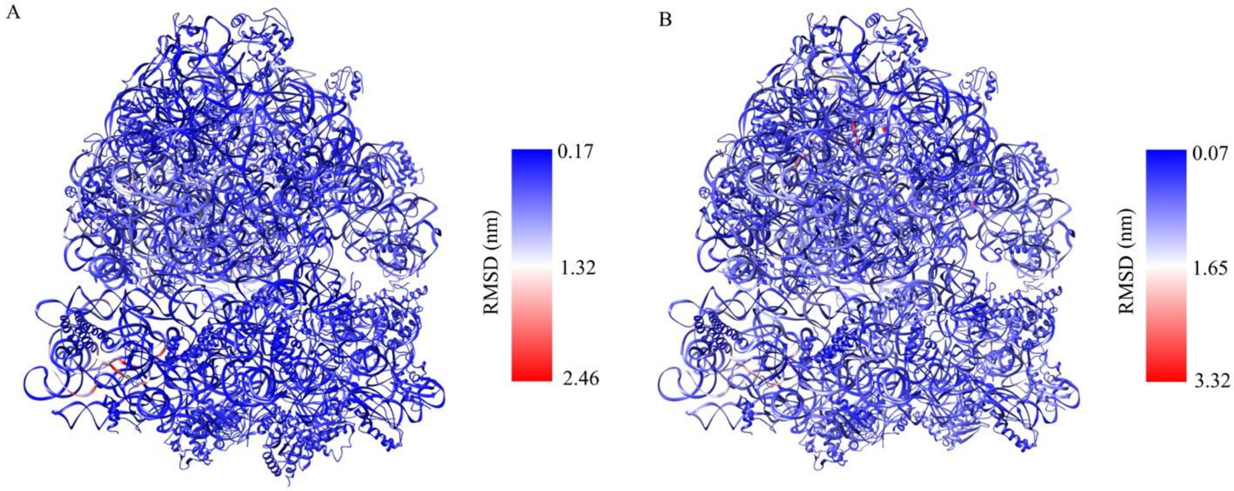


**Figure S6. RMSD comparison.** Comparison of RMSD of rRNA and its associated proteins between MRE600 and (A) wB11 or (B) nB11.

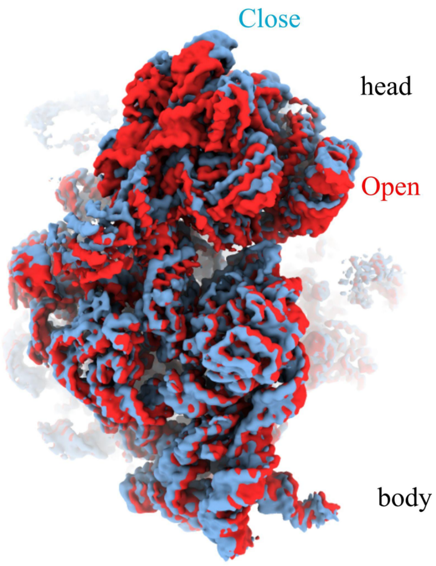


**Figure S7. Latch-opening/closing.** The beginning and end states of the first two principal components are compared to show the "nodding" latch-opening/closing head movement of nB11.

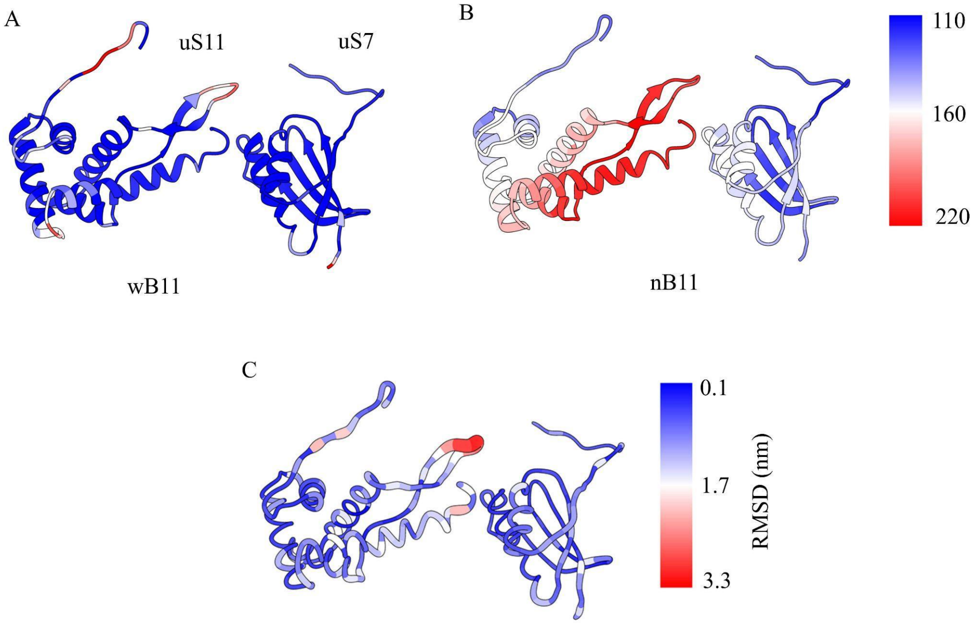


**Figure S8. B-factor and RMSD comparison.** B-factors of two proteins uS11 and uS7 of (A) wB11 and (B) nB11. (C) is the comparison of RMSD of two proteins uS11 and uS7 between wB11 and nB11.

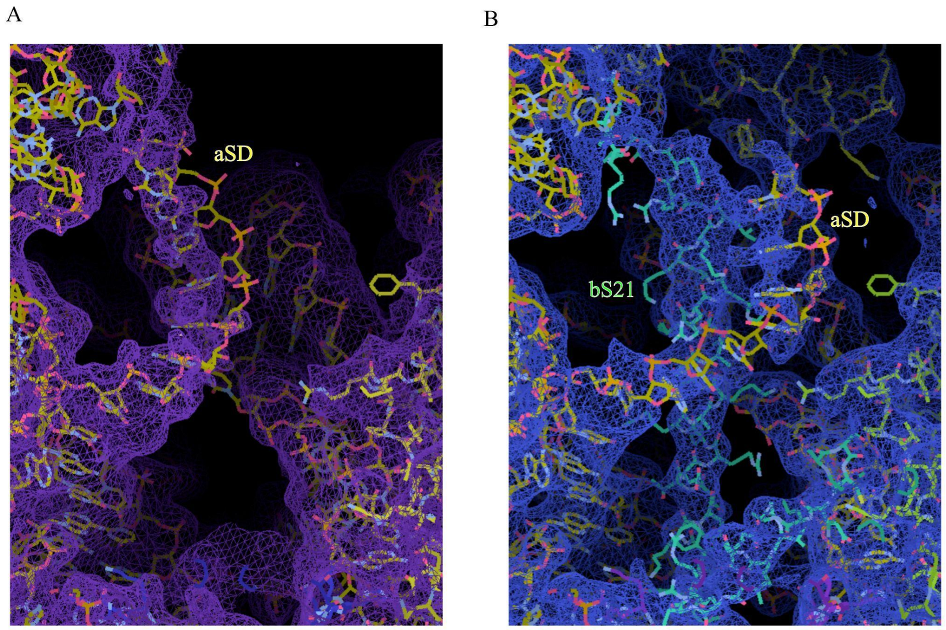


**Figure S9. Displacement of aSD.** Densities of aSD in (A) nB11 and (B) wB11 showing the displacement of aSD in the absence of bS21.

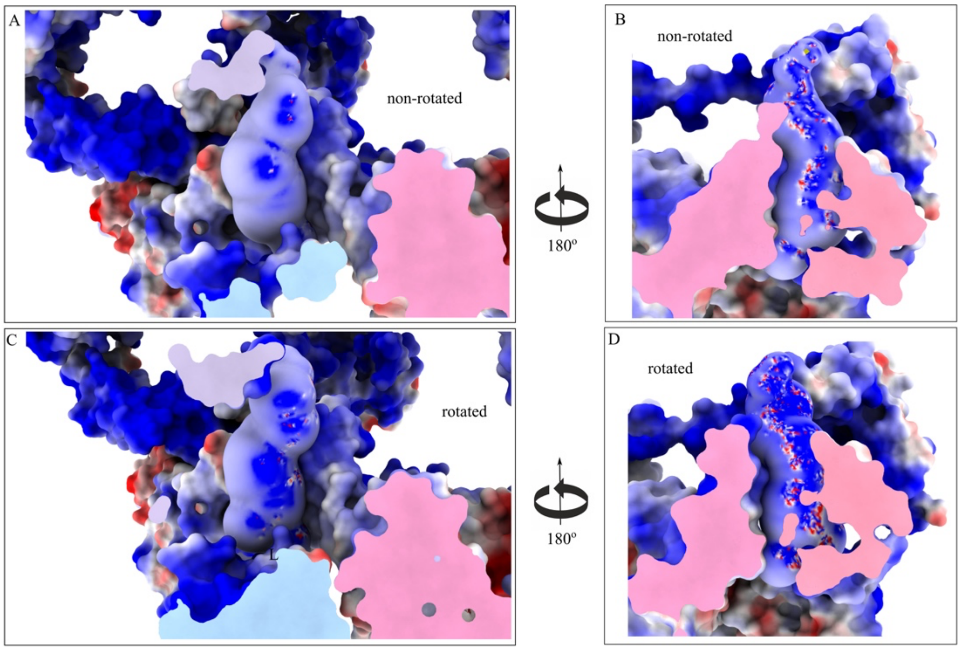


**Figure S10. Charge distribution in the lmRNA entrance channel.** (A,B) explicit charge distribution in the channel for class C3m; (C,D) explicit charge distribution in the channel for class C5m. Surface representations of the channel insides were calculated using MOLEonline (https://mole.upol.cz/).

| Map and Model component | MR600 | wB11 | nB11 |
|---|---|---|---|
| Map resolution (Å)<br>FSC threshold 0.143 (Å) | 3.19 | 3.2 | 3.3 |
| Map sharpening *B* factor (Å$^2$) | -76.43 | -87.01 | -89.81 |
| Model composition (residues) | | | |
| Protein | 5496 | 5496 | 5227 |
| Nucleotide | 4648 | 4648 | 4648 |
| *B* factors (Å$^2$) | | | |
| RNA | 49.06 | 56.96 | 66.28 |
| protein | 42.94 | 56.68 | 64.88 |
| R.m.s. deviations from ideal values | | | |
| Bond (Å) | 0.010 | 0.004 | 0.008 |
| Angle (°) | 1.026 | 0. 848 | 0.971 |
| Molprobity score | 1.55 | 1.55 | 1.55 |
| Clash score | 4.84 | 4.67 | 4.87 |
| Ramachandran plot (%) | | | |
| Favored (%) | 95.66 | 95.57 | 95.79 |
| Allowed (%) | 3.04 | 4.32 | 4.21 |
| Outliers (%) | 0.02 | 0.11 | 0.00 |
| Rotamer outliers (%) | 0.93 | 0.56 | 0.89 |
| CB outliers (%) | 0.02 | 0.00 | 0.02 |
| RNA validation | | | |
| Good_Sugar Puckers(%) | 99.99 | 99.99 | 99.83 |
| Good_backbone_conformation (%) | 99.81 | 99.99 | 99.99 |
| Average suiteness | 0.501 | 0.516 | 0.507 |

**Table S1: Model refinement statistics.**
